# Mitochondrial survivin reduces oxidative phosphorylation in cancer cells by inhibiting mitophagy

**DOI:** 10.1101/2020.04.21.052662

**Authors:** Amelia R. Townley, Sally P. Wheatley

## Abstract

Survivin is a cancer-associated protein that is pivotal for cellular life and death: it is an essential mitotic protein and an inhibitor of apoptosis. In cancer cells, a small pool of survivin localises to the mitochondria, the function of which remains to be elucidated. Here, we report that mitochondrial survivin inhibits the selective form of autophagy, called “mitophagy”, causing an accumulation of respiratory defective mitochondria. Mechanistically the data reveal that survivin prevents recruitment of the E3-ubiquitin ligase Parkin to mitochondria and their subsequent recognition by the autophagosome. The data also demonstrate that, as a consequence of this blockade, cells expressing high levels of survivin have an increased dependency on anaerobic glycolysis. As these effects were found exclusively in cancer cells they suggest that the primary act of mitochondrial survivin is to force cells to implement the “Warburg Effect” by inhibiting mitochondrial turnover.

## Introduction

Survivin is a protein at the interface of cellular life and death, as it guides mitosis and inhibits apoptosis (Wheatley and Altieri, 2019). It is over-expressed in cancer and is associated with poor patient prognosis (Ambrosini et al. 1997; Escuín & Rosell 1999, reviewed in Jaiswal et al. 2015). Its abundance correlates directly to chemotherapy resistance highlighting it as a potential anti-cancer target (Morrison *et al*., 2012). Survivin localises in several distinct pools (Fortugno *et al*., 2002). During mitosis and as part of the chromosomal passenger complex, it directs chromosome congression and segregation, as well as cytokinesis. When present in interphase, its predominantly cytosolic localisation is key to its anti-apoptotic function. The focus of this paper is the mitochondrial pool of survivin, which is found only in cancer cells (Dohi *et al*., 2004). What survivin does when resident in the mitochondria is a matter of ongoing debate (Hagenbuchner *et al*., 2013; Rivadeneira *et al*., 2015), although early evidence suggested that it may be a store of survivin with greater anti-apoptotic potential than the cytosolic pool “primed” in readiness to respond to pro-apoptotic signals (Dohi, Xia and Altieri, 2007).

Malignant transformation requires cellular changes that enable unrestricted proliferation and circumvention of programmes of cell death, such as apoptosis and autophagy. For decades the mitochondrion has been seen as a by-stander of malignant transformation, gaining inactivating mutations from various sources that switch cellular metabolism from oxidative phosphorylation (OXPHOS) to glycolysis (Warburg, 1956). More recently it has begun to be appreciated that mitochondria actively participate in driving tumour progression and malignant transformation (Chatterjee, Dasgupta and Sidransky, 2011; Yadav and Chandra, 2013; van Gisbergen *et al*., 2015)

Mitochondrial homeostasis, including their quality and length is maintained by the dynamic processes of fusion and fission, which are controlled by factors within the mitochondria and in the cytosol (Westermann, 2010a; East and Campanella, 2016). In healthy cells, the balance between fusion and fission is tightly controlled to allow for the timely removal of non-functional mitochondria without affecting respiration (Nunnari *et al*., 1997). In cancerous cells, this process is commonly deregulated resulting in the gradual accumulation of mitochondrial DNA (mtDNA) mutations that eventually trigger the loss of the respiratory apparatus (Balaban, Nemoto and Finkel, 2005; Porporato *et al*., 2017), causing a reduction in OXPHOS, and greater dependency on glycolysis, hence evoking the “Warburg Effect” (Merz and Westermann, 2009). In normal proliferating cells, fusion is important for the maintenance of healthy mitochondria as it can rescue damaged mitochondria by mixing their contents with healthy mitochondria (Westermann, 2010b). Opposing fusion is fission, a process that generates mitochondrial fragments. In mitosis fission occurs to ensure that the organelle is correctly inherited (Twig, Hyde and Shirihai, 2008), but it is also necessary to maintain mitochondrial homeostasis as it precedes the removal of defective mitochondria by the selective form of autophagy, called “mitophagy” (Twig and Shirihai, 2011; Redmann *et al*., 2014). Together, with mitochondrial biogenesis, mitophagy is a quality control mechanism that determines mitochondrial mass (Jornayvaz and Shulman, 2010).

Autophagy delivers defective organelles to autophagosomes, which fuse with lysosomes to degrade and recycle their constituents (Youle and Narendra, 2011). Organelles destined for recycling can be delivered to the autophagosome by two distinct pathways, either ubiquitin-dependent or ubiquitin-independent (Zaffagnini and Martens, 2016). The ubiquitin-dependent pathway requires ubiquitination of defunct organelles by E3-ubiquitin ligases and their subsequent recognition by ubiquitin binding proteins, which enables extension of the pre-autophagosomal (phagophore) membrane and their complete engulfment into the autophagosome (Shaid *et al*., 2013). Mitochondrial recycling relies upon the action of PTEN-induced kinase 1 (PINK) and the E3-ubiquitin ligase Parkin (East and Campanella, 2016). Alterations to either of these processes result in the accumulation of defunct, metabolically inactive mitochondria. Reactive oxygen species (ROS), which are a principle cause of mitochondrial DNA (mtDNA) damage, are a by-product of OXPHOS. If mitochondria harbouring mtDNA lesions are not removed in a timely manner they can promote tumourigenesis (Ott *et al*., 2007). Cancer cells typically circumvent the damage by using glycolysis to generate ATP. Even though it is a less efficient means of ATP production, one of its major advantages is that it does not produce ROS and thus does not cause mtDNA mutations. Mitochondrial quality control is a relatively unexplored aspect of cancer, but given the accumulating evidence that alterations in mitophagy can bestow chemotherapy resistance (van Gisbergen *et al*., 2015), understanding its contribution to the diseased state may reveal an Achilles’ heel of cancer (Hagenbuchner *et al*., 2016)

Here we test the hypothesis that survivin regulates mitochondrial homeostasis and respiratory dependence in cancer cells. We report that in cancer cells, survivin increases mitochondrial mass and reduces mtDNA quality by inhibiting mitophagy. We propose that due to this accumulated load of respiratory inactive mitochondria, cancer cells with high expression of survivin undertake the Warburg transition and become reliant on glycolysis for survival.

## Results

### Survivin is found in the mitochondria of transformed cells

It has previously been reported that a sub-population of survivin localises to the mitochondria in cancer cells. To verify that this was the case in the cells being examined here, we carried out subcellular fractionation to enrich for mitochondria in two cancerous lines: HeLa (cervical cancer) and U2OS (osteosarcoma), and in normal fibroblasts (MRC5), see Figure 1A. All lines were engineered to ectopically express survivin, C-terminally tagged with GFP or GFP alone (control). As indicated by enrichment of VDAC, and minimal contamination of tubulin (cytosolic marker) and histone H3 (nuclear marker), while GFP (control) was excluded from mitochondria in all lines, survivin-GFP was present in the mitochondria of HeLa and U2OS, but not MRC5 cells. These data corroborate previous work (Dohi *et al*., 2004), and further demonstrate that this is also the case when survivin is present at high levels through ectopic expression.

**Figure 1.**
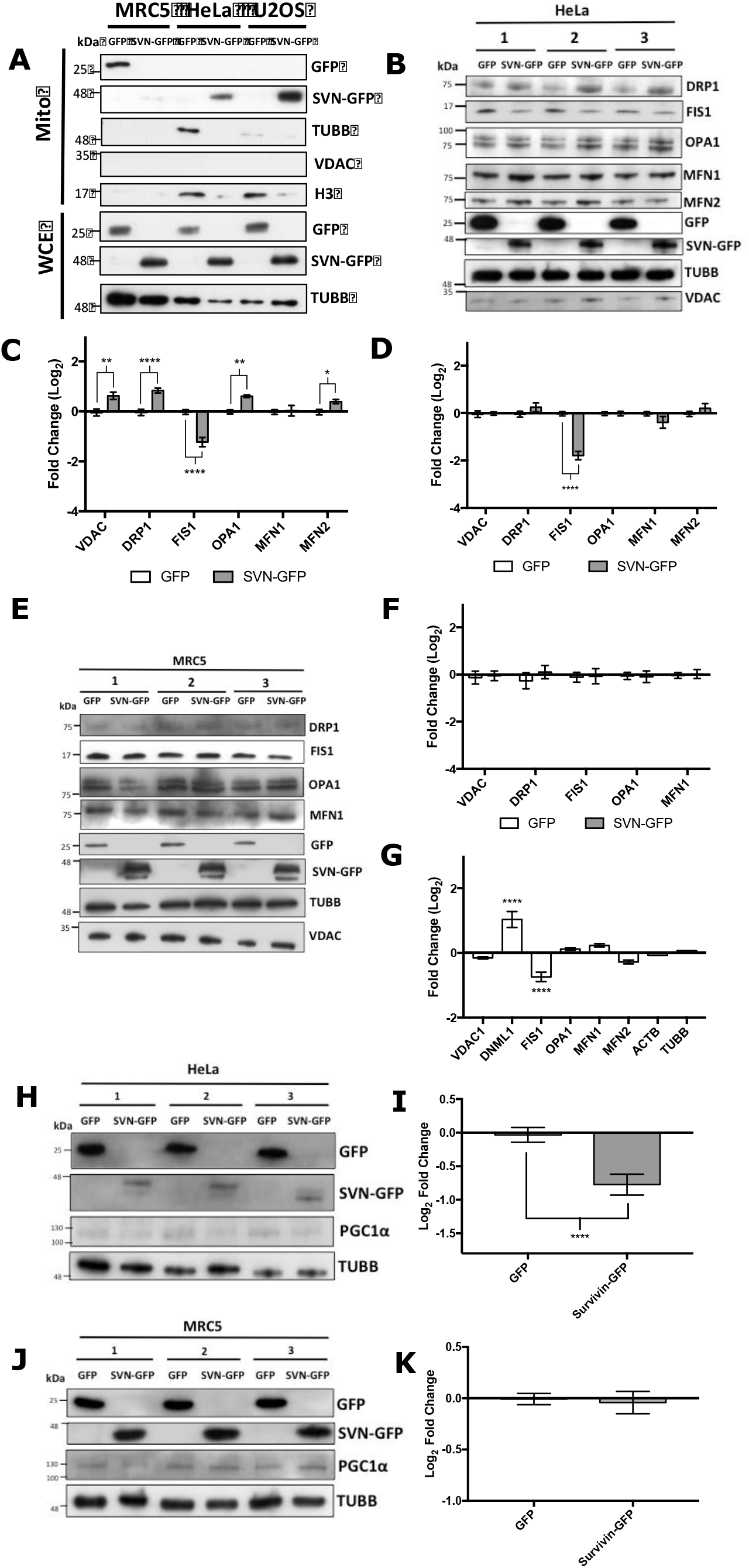
Analysis of mitochondrial proteins in cancer and normal cells. **(A)** Immunoblot of the mitochondria enriched fractions (Mito) from GFP and survivin-GFP (SVN-GFP) overexpressing MRC5, HeLa and U2OS cells. Whole cell extracts (WCE) are included in the lower panel to indicate equality in expression of each ectopic protein. VDAC is a mitochondrial (OMM) marker; Histone 3 indicates any nuclear contamination, and tubulin serves both an indicator of cytoplasmic contamination (Mito) and a loading control (WCE). **(B-F)** Immunoblot analysis of mitochondrial protein expression in WCEs from HeLa sublines indicated. Compared to GFP controls SVN-GFP expressing cells had increased VDAC, OPA1 and MFN-2 expression and decreased FIS1 expression. (**C)** Semi-quantitative analysis of (B). Pixel intensity was normalised to tubulin and compared to the GFP control. A two-way ANOVA was used to test significance, p<0.05 (*), p<0.01 (**), p<0.001 (***), p<0.0001(****). **(D)** Semi-quantitative analysis of immunoblot in (B) normalised to VDAC, statistically analysed using a two-way ANOVA test, p<0.0001(****). Changes in FIS1 remained significant. **(E)** Experiment described in (A) carried out in MRC5 cells. **(F)** Semi-quantitative analysis of immunoblot in (E) normalised to tubulin. Pixel intensity quantified as described above (B). Two-way ANOVA shows no statistical differences in protein expression caused by SVN-GFP expression. All error bars indicate +/- SEM, N=3 (triplicate). **(G)** qPCR analysis of mRNA to the proteins analysed in B-F expression in HeLa cells over-expressing GFP or SVN-GFP. VDAC mRNA remains constant, DRP1 and FIS1 expression are significantly increased and decreased respectively. **(H)** WCE’s from HeLa or **(J)** MRC5 cells over-expressing GFP or SVN-GFP and analysed by immunoblotting. Membranes were probed for PGC1α as a mitochondrial biogenesis marker, GFP to check cell line expression and tubulin as a loading control, N=3 (with internal triplicates). **(I)** and **(K)** Semi-quantification of immunoblots in **(H)** and **(J)** respectively, presented as fold change (Log2 scale) compared to the GFP control (TWO-way ANOVA P-value **** p<0.0001). SVN-GFP expression in HeLa cells reduces the expression of PGC1α but causes no alterations in MRC5 cells.

### Manipulating survivin expression alters mitochondrial mass and affects expression of fusion and fission proteins in cancer cells, but not proteins required for mitochondrial biogenesis

To determine whether mitochondrial survivin can influence expression of mitochondrial proteins, including those responsible for mitochondrial dynamics, we analysed whole cell extracts (WCE) of HeLa cells expressing GFP or survivin-GFP by immunoblotting (Figure 1B). Blots were probed for the fission proteins DRP1 and FIS1, the fusion proteins, MFN1, MFN2 (OMM), and OPA1 (IMM); mitochondrial mass was assessed using anti-VDAC. Semi-quantitative analysis of these blots (Figures 1C) was calculated by normalising the band intensity of the protein of interest against the anti-tubulin loading control and presented as fold change of survivin-GFP compared with GFP. This analysis showed that VDAC levels increased significantly when survivin was expressed. In addition to this, DRP1, MFN-2 and OPA1 expression also increased, while the fission receptor protein, FIS1, decreased. Given that total mitomass increased, we recalculated the fold change normalised to VDAC. This revealed that only the change in FIS1 occurred independently of mitomass (Figure 1D). In contrast to the results in HeLa cells, none of these alterations were observed in normal MRC5 fibroblasts expressing survivin-GFP (Figure 1E & F). To determine whether changes in expression of these proteins could be attributed to changes at the transcriptional level, quantitative PCR (qPCR) was performed on extracts from HeLa cells and fold change in expression between survivin-GFP and GFP controls plotted. No change in VDAC mRNA was observed but there was a statistically significant increase in DRP1 mRNA (encoded by DNML-1), and a decrease in FIS1 mRNA (Figure 1G), which could account for the changes observed at the protein level.

To clarify whether the increase in mitomass was caused by elevated mitochondrial biogenesis, HeLa and MRC5 WCEs expressing GFP or survivin-GFP were run on an SDS-page gel and immunoblotted with the biogenesis marker PGC1-α (Figure 1H and J). Semi-quantitative analysis showed that protein expression was decreased in HeLa cells (Figure 1I) and not altered in MRC5 cells (Figure 1K). Thus, we conclude that the observed increase in mitomass is not due to increased mitochondrial biogenesis.

Having established that survivin overexpression increases mitomass and decreases FIS1 levels, we next asked whether depleting it would have the opposite effect. To investigate this survivin-specific siRNA was performed (48h) on HeLa and MRC5 cells (Figure S1). WCEs were separated by SDS-PAGE, transferred to nitrocellulose and probed for mitochondrial proteins as described above. Semi-quantitative analysis of blots was used to determine the fold change of each protein in siRNA treated versus untreated cells and normalised to tubulin. Survivin depletion caused a significant reduction in VDAC and MFN1 expression and no change to FIS1 in HeLa cells (Figure S1A and C). By contrast none of these effects were observed in MRC5 cells (Figure S1B and D). These data complement the overexpression data, and collectively prove that changes in survivin expression alter mitochondrial mass in cancerous but not in normal cells.

### Survivin increases mtDNA copy number but mtDNA integrity is compromised

To confirm that survivin expression increases mitochondrial mass, we quantified mitochondrial DNA (mtDNA) copy number. Genomic DNA was extracted from HeLa or MRC5 cells expressing either GFP or survivin-GFP, or after siRNA depletion. qPCR was used to determine the abundance of the mtDNA encoded tRNA(LEU) gene, which was then compared to the stably expressed nuclear reference genes ACTB (actin) and TUBB (tubulin). Fold change of RNA between cells expressing survivin-GFP and GFP was then calculated and presented on a Log2 scale. Survivin overexpression increased mtDNA tRNA(LEU) gene expression in HeLa cells (Figure 2A), and decreased it in MRC5 cells (Figure 2B). Thus, by an independent method, these data concur that mitomass is elevated by survivin expression in cancer cells.

**Figure 2.**
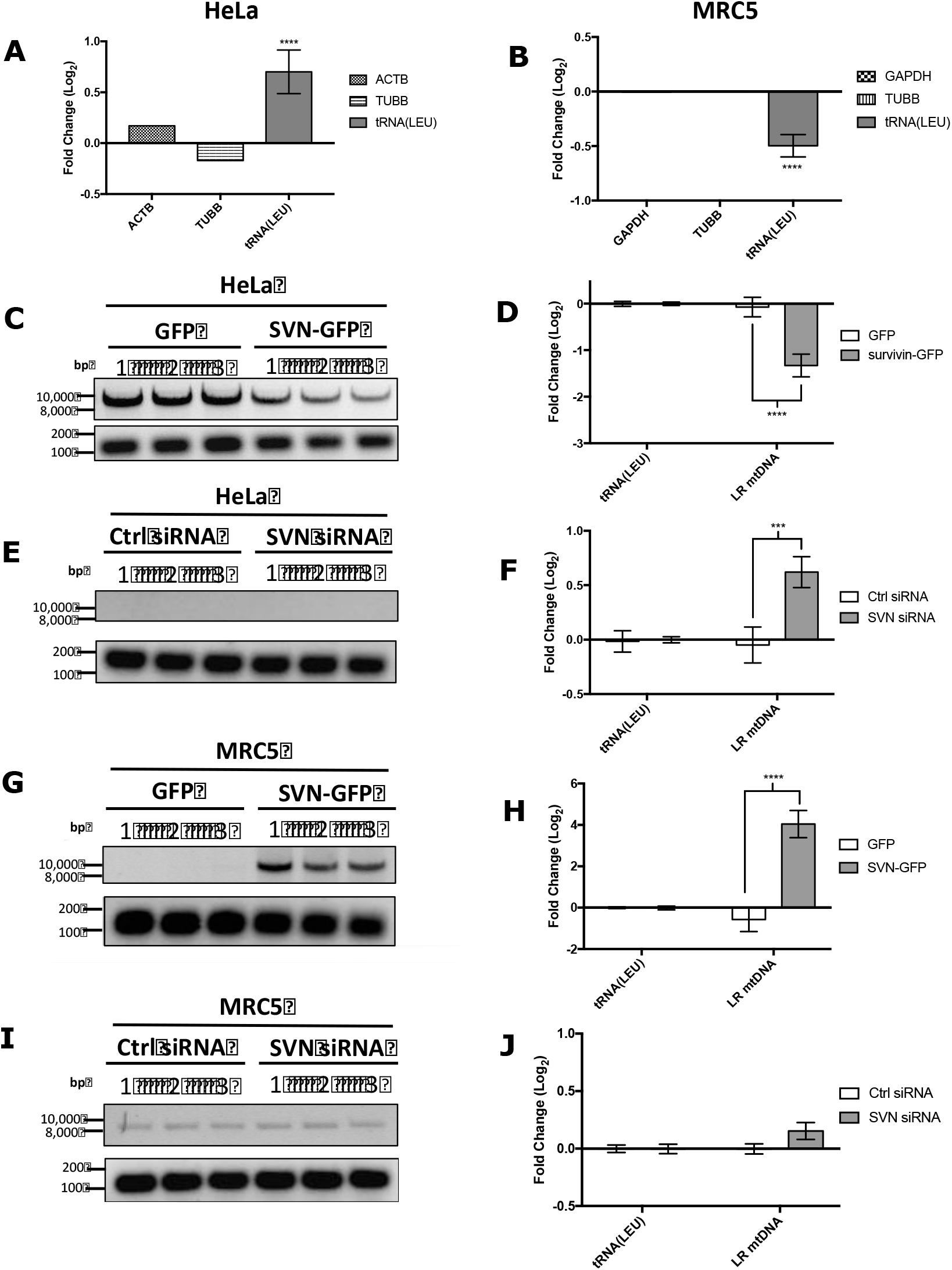
mtDNA copy number increases but its quality decreases in HeLa cells expressing survivin-GFP. mtDNA copy number was determined by quantitative PCR on genomic DNA extracted from **(A)** HeLa and **(B)** MRC5 cells expressing GFP or SVN-GFP. The mitochondrial encoded tRNA(LEU) gene was quantified and compared to two stably expressed nuclear reference genes, tubulin (TUBB) and actin (ACTB) or GAPDH. tRNA(LEU) was increased and decreased in SVN-GFP expressing HeLas and MRC5 cells respectively (p<0.0001). **(C, E, G** and **I)** Long read (LR: 9 kb) and short read (SR: 150 bp) PCR products derived from genomic DNA extracted from **(C)** HeLa cells and **(G)** MRC5 cells expressing GFP or SVN-GFP, or survivin knockdown with siRNA (SVN siRNA) **(E)** and **(I)**. LR fragments were reduced in survivin-GFP HeLas (p<0.0001) but increased after SVN siRNA (p=0.0008). In comparison, in MRC5 cells the 9 kb product increased (p<0.0001), and no change was observed after survivin depletion. **(D, F, H** and **J)** Semi-quantitative analysis using Image J and Prism software of the 9 kb band intensity normalised to the 150 bp product. Statistical significance was determined using a two-way ANOVA test, p<0.001 = ****, points show mean +/- SEM N=3 (triplicate).

Next, we examined the quality of the mtDNA using a PCR-based lesion frequency assay. Briefly, genomic DNA was prepared from cells as described for Figure 2A and B and PCR reactions carried out to produce either a ‘long read’ (9kb) or a ‘short read’ (150 bp) product. In this assay if mtDNA is intact both products will be generated, however, if the DNA polymerase encounters lesions, it will stall and less 9kb product will form. The 150 bp product is used as a loading control. As shown in Figure 2C and D, when normalised to the 150 bp band and analysed semi-quantitatively, fewer 9kb products were generated in HeLa cells expressing survivin, compared to the GFP control, indicating their mtDNA had more lesions. Conversely survivin depletion increased the 9kb product suggesting that in these cells the mtDNA was intact (Figures 2E and F). In contrast, in MRC5 cells survivin expression actually increased the production of the 9kb product (Figures 2G and H), whereas its depletion had no effect (Figure 2I and J). Taken together these data suggest that survivin increases mtDNA copy number but reduces its quality specifically in cancer cells.

### Survivin expression reduces oxidative phosphorylation in cancer cells

As mitochondrial quality was impaired, we next asked whether survivin expression also affected respiration. To address this a resazurin assay was carried out at 0.5h intervals over 4h on mitochondria isolated from HeLa, U2OS or MRC5 cells (Figure 3). Both HeLa and U2OS cells expressing survivin-GFP showed a significant decrease in resorufin fluorescence compared to GFP expressing cells (Figure 3A & C). To ensure the observed alterations were due to changes in OXPHOS and not the mitochondrial TCA cycle, we also determined the response of each line to the complex V inhibitor, oligomycin (Figures 3B, D and F). Mitochondria isolated from both HeLa and U2OS cells expressing GFP were more sensitive to oligomycin than those from survivin-GFP cells (Figure 3B & D), suggesting that survivin significantly reduced OXPHOS in cancer cells. In contrast, no difference was seen in the ability of MRC5-derived mitochondria to metabolise resazurin (Figure 3E and S2). In addition, MRC5 cells expressing survivin-GFP tagged with a *bona fide* mitochondrial targeting sequence from cytochrome c oxidase subunit VIIIA, forcing survivin to be mitochondrial, reduced mitochondrial metabolism in a manner seen as in HeLa and U2OS cells (Figure 3F). From these data we conclude that survivin specifically inhibits reduction reactions when present within the mitochondria.

**Figure 3.**
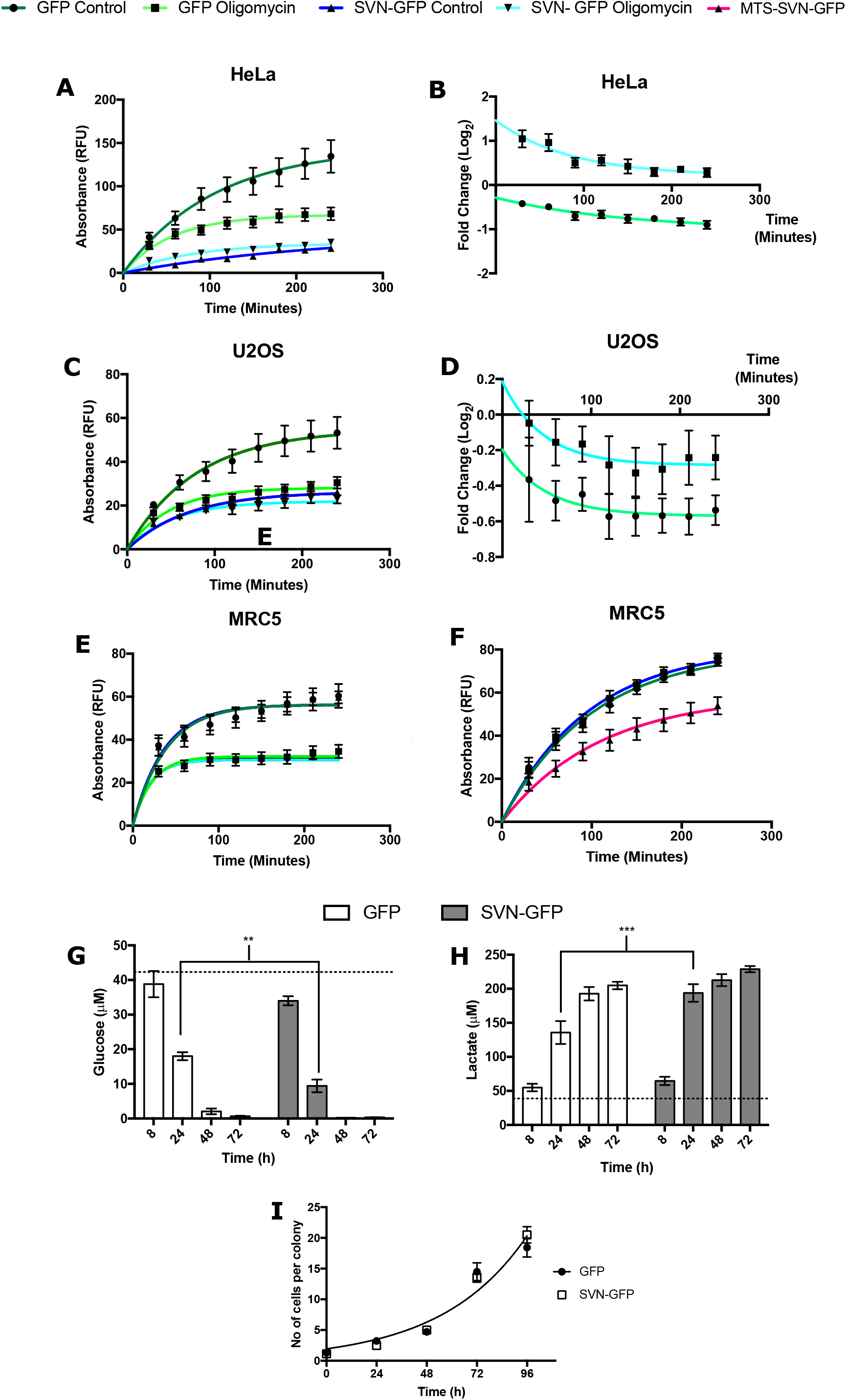
Analysis of mitochondrial respiration, response to oligomycin and lactate production. Mitochondria were isolated from **(A)** HeLa, **(C)** U2OS or **(E)** MRC5 cells expressing GFP or SVN-GFP and plated in resazurin medium with or without oligomycin and metabolism assessed by absorbance, measured in relative fluorescence units (RFU) every 30 minutes for 4 h. Using non-linear regression analysis there was no line of best fit matching any pair of data sets in (**A**) and (**C**) analysis (P-value <0.0001), but a single curve fitted all datasets for (**E**); GFP, SVN-GFP, GFP + oligomycin and SVN-GFP + oligomycin (p values 0.9808 and 0.4123 respectively). Mean +/- SEM, N=3 is plotted (internally in triplicate). **(B)** and **(D)** Mitochondria from HeLa and U2OS display a greater fold change in resorufin fluorescence with oligomycin treatment compared to SVN-GFP mitochondria. Non-linear regression analysis shows no curve fits both HeLa/U2OS GFP and SVN-GFP oligomycin treatment data points (p value <0.0001). **(F)** Mitochondria isolated from MRC5 cells expressing GFP, SVN-GFP or MTS-SVN-GFP and analysed as mentioned previously. Non-linear regression analysis demonstrates that MTS-SVN-GFP expression reduces resorufin fluorescence in comparison to GFP or SVN-GFP (P-value <0.0001), Mean +/- SEM, N=3 (internally in triplicate). **(G)** Glucose consumption of 10,000 cells was measured over 72 h using a glucose-Glo assay. Two-way ANOVA test at 24h was used to determine significance (P-0.0026). Regression analysis to analyse difference in rate of change, F test proves significant difference (p value - 0.0176). **(H)** Lactate production was measured as for **(G)** using a lactate-Glo assay. Two-way ANOVA test at 24h was used to determine significance (P value - 0.0005). Regression analysis and F-test proves significant differences (p value - 0.0029). Dotted line shows glucose or lactate concentration at 0h. Mean +/- SEM is presented, N=2 (internally in triplicate). **(I)** GFP and SVN-GFP expressing HeLa cells grow at the same rate. 200 HeLa cells were seeded and the number of cells per colony counted over 72 h. Mean +/- SEM is presented, N=3.

### Survivin expression increases glucose consumption and lactate production

As survivin overexpression was found to reduce OXPHOS, we next asked whether cancer cells compensated for this reduction by increasing anaerobic respiration. To test this, we used luciferase-based assays to measure glucose consumption and lactate production. Survivin-GFP HeLa cells had a significantly lower glucose concentration (Figure 3G) and higher rate of lactate production (Figure 3H) compared to GFP expressing cells 2h post-seeding. Moreover, the rates at which glucose was consumed and lactate concentration rose was significantly higher than those observed in GFP cells. As both cell lines grew at the same rate (Figure 3I), we conclude that the differences observed were due primarily to metabolic adjustments and not differences in proliferation.

### Survivin reduces the sensitivity of cancer cells to chloroquine

Based on the observations thus far we hypothesised that survivin inhibits the removal of defective mitochondria by the selective autophagic process of “mitophagy”. To test this, cells over-expressing GFP or suvivin-GFP were exposed to the autophagy inhibitor chloroquine (CQ) for 16 h, the mitochondria isolated and a resazurin assay performed. As shown in Figures 4A-D, CQ treatment (50 and 150 μM) reduced metabolism of mitochondria isolated from GFP-expressing cancer cells, which is consistent with a block on the removal of defunct mitochondria. However, mitochondria from survivin-GFP cells showed no reduction in metabolism, which even slightly increased in response to CQ. Consistent with the lack of survivin in the mitochondria of normal cells, those isolated from MRC5 GFP or survivin-GFP cells responded similarly to CQ (Figure 4E and F). Taken together these data suggest that metabolically compromised mitochondria accumulate when survivin is expressed because it inhibits mitophagy.

**Figure 4.**
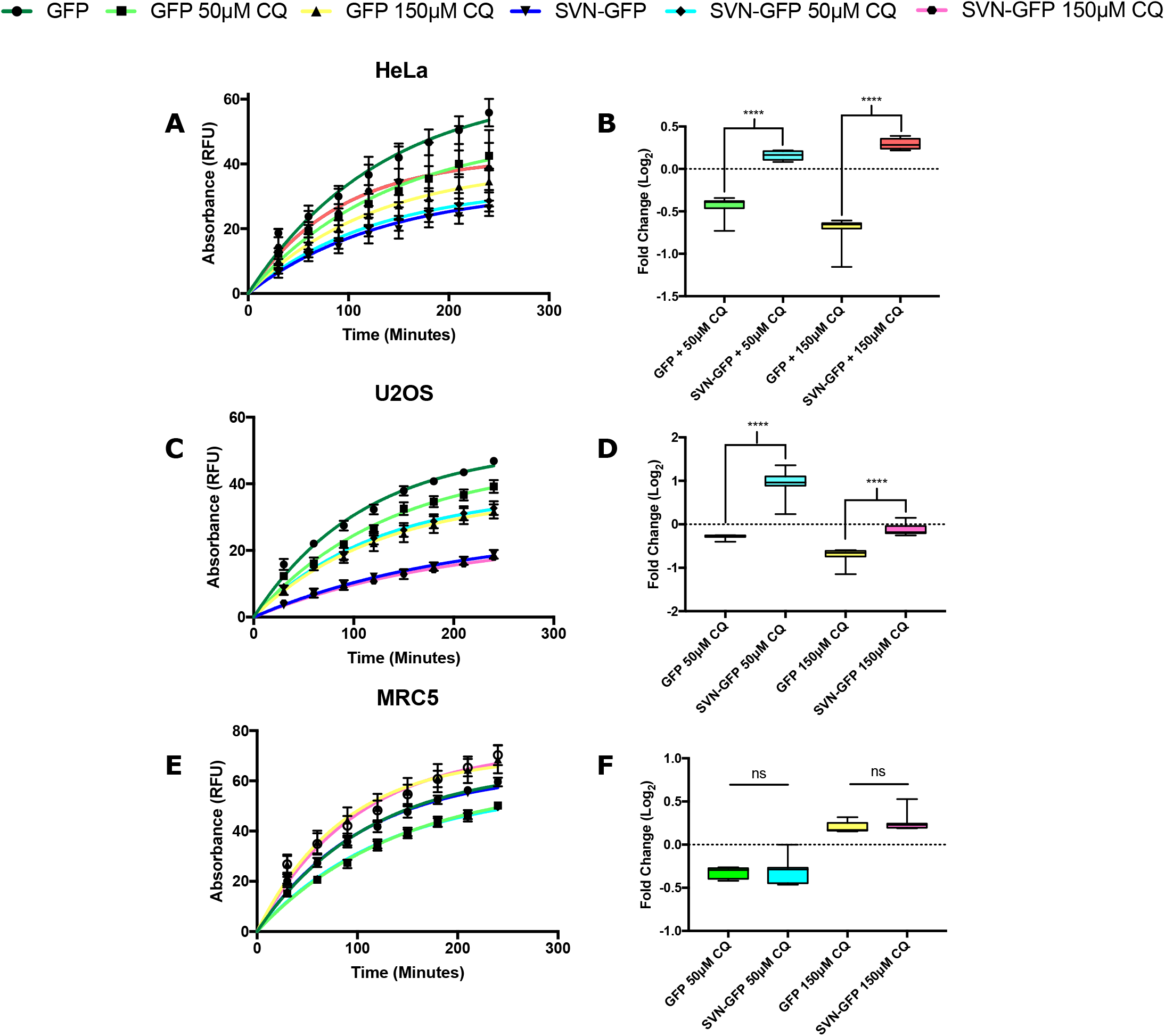
Effect of chloroquine (CQ) treatment on mitochondrial respiration. Mitochondria isolated from **(A** and **B)** HeLa and **(C** and **D)** U2OS cells expressing GFP or SVN-GFP were plated with the addition of resazurin +/- 50 μM or 150 μM CQ and fluorescence absorption measured read every 30 minutes for 4 h. Non-linear regression shows no line fits any pair of data sets in control GFP expressing HeLa or U2OS mitochondria, whereas a statistical increase is seen in response to each CQ treatment in HeLa and U2OS cells expressing SVN-GFP, mirrored in fold change bar chart analysis. **(E** and **F)** Mitochondria isolated from MRC5 cells expressing GFP or SVN-GFP display the same fold change in resorufin fluorescence post-CQ treatment. Non-linear regression analysis shows line fits both MRC5 GFP and SVN-GFP 50 μM or 150 μM CQ. All points indicate mean +/- SEM, N=3 (triplicate). No statistical significance is observed in the fold change between GFP and SVN-GFP 50 μM or 150 μM CQ treatments.

### Survivin-depletion causes mitochondrial fragmentation in cancer cells

Next, we used a combination of fluorescent mitochondrial stains and live imaging to observe the total mitochondrial network in HeLa cells transiently overexpressing GFP/RFP or survivin-GFP/survivin-RFP, or depleted of survivin. In HeLa cells mitochondrial morphology was unaffected by ectopic expression of survivin (Figures 5A and B, S3), however, consistent with the immunoblotting and qPCR data, when total pixel intensity was measured it was apparent that mitomass was increased when survivin-RFP was expressed (Figure 5C). Conversely, its depletion caused mitochondrial fragmentation (Figures 5E and F, S4A). We then monitored the mitochondrial membrane potential using TMRE or MitoTracker Far Red; two fluorescent molecules that only highlight polarised mitochondria. The average signal intensity was quantified and normalised to the average intensity of MitoTracker Green to account for mitochondrial mass (TMRE/MTFR:MTG). In HeLa cells, there was a slight reduction in OMM potential in the presence of survivin-GFP, but using this particular combination of dyes, (which was necessary for imaging purposes) it was not deemed significant (Figure 5D). In the converse experiment survivin depletion reduced the mitochondrial membrane potential (Figure 5G), and in MRC5 cells no alterations to mitochondrial morphology (Figure 5H) or membrane potential were observed after survivin ablation (Figure 5I, S4B). Thus, we conclude that specifically in cancer cells, loss of endogenous survivin increases mitochondrial fission and can depolarise the OMM.

**Figure 5.**
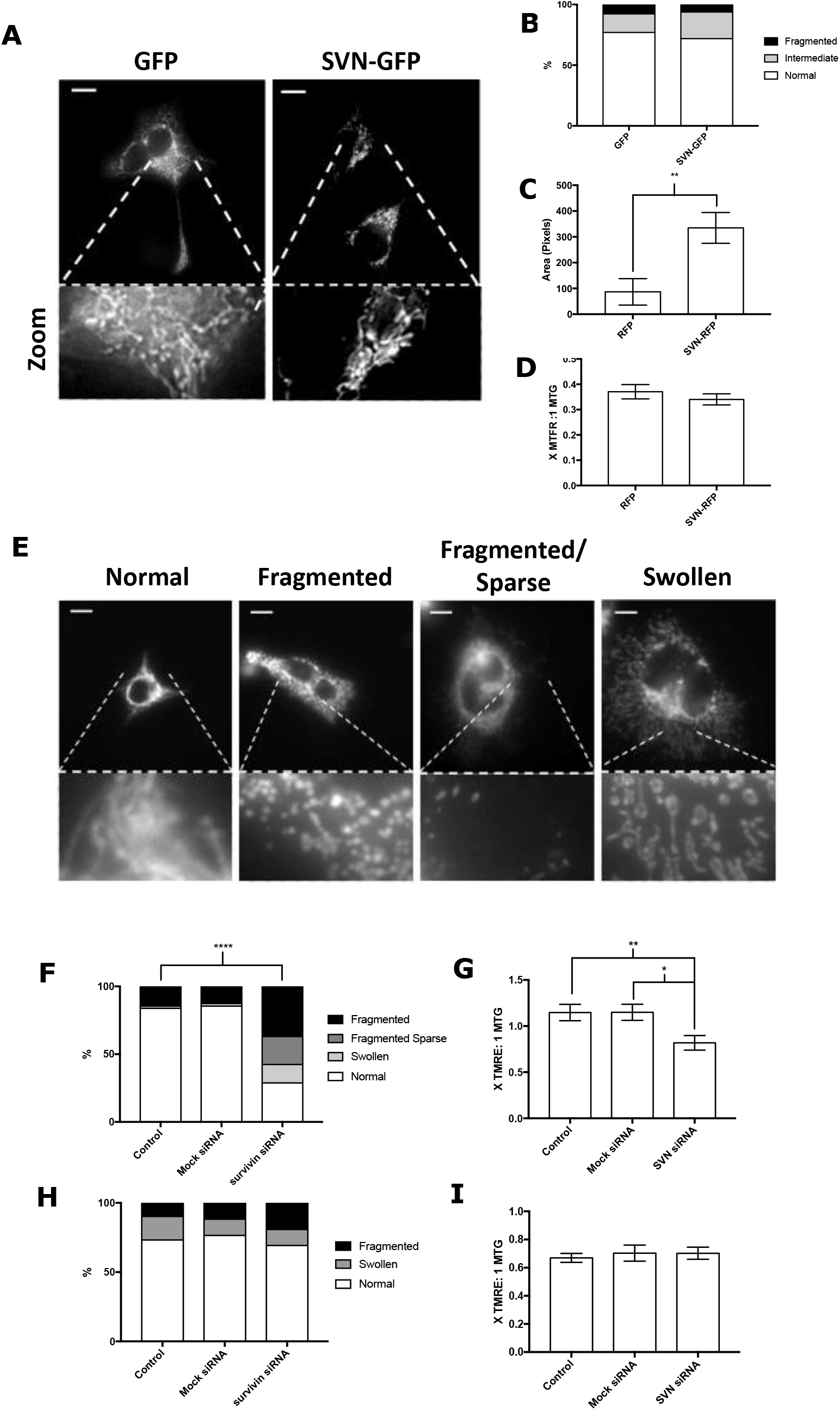
Survivin depletion increases mitochondrial fragmentation and decreases mitochondrial membrane potential in cancer cells. **(A** and **B)** HeLa cells expressing GFP or SVN-GFP were stained with 250 nM MitoTracker Red FM and imaged live: 3 mitochondrial phenotypes were observed; normal, intermediate or fragmented. Chi-squared test indicated a similar mitochondrial distribution in both lines regardless of survivin status, N=3, DF=2. Full galleries are shown in supplementary Figures S2 and S3. **(C** and **D)** Survivin overexpression does not alter mitochondrial membrane potential in HeLa cells but does increase mitochondrial pixel area. HeLa cells transiently expressing RFP or SVN-RFP were stained with MitoTracker Green (MTG) or MitoTracker Far Red (MTFR) and imaged live. Images were thresholded using Fiji software, mean pixel intensity and area was quantified and MTFR signal normalised to MTG. (TWO-way ANOVA, P-value ****p<0.0001). **(E** and **F)** HeLa cells were treated with control or survivin-specific siRNA, stained with MTG. Four mitochondrial phenotypes were observed; normal, fragmented, fragmented sparse and swollen. SVN-siRNA increased mitochondrial fragmentation (Chi-squared test, N = 3, DF=3). **(G)** Experiment described in (**F**) but HeLa cells were stained with MTG and TMRE. One-Way ANOVA indicated a statistically significant reduction in TMRE signal compared after SVN-siRNA, after normalisation to MTG. **(H** and **I)** MRC5 cells were treated and imaged as described in (**F** and **G**): neither the mitochondrial network nor its membrane potential was unaffected by survivin-depletion (Chi-squared test, N = 3, DF=3).

### Survivin does not inhibit mitophagic steps preceding mitochondrial translocation of Parkin

As experiments described thus far suggested that survivin inhibits mitophagy, our final aim was to determine where survivin was operating in the mitophagic pathway. To address this HeLa cells expressing GFP/survivin-GFP or RFP/survivin-RFP were treated with FCCP (10 μM, 6h) to depolarise the mitochondria and stimulate mitophagy. They were then stained with MitoTracker Red to visualise the mitochondrial network, and MitoTracker Green/MitoTracker Far Red to determine membrane polarisation respectively. As shown in Figure 6A and B, FCCP caused mitochondrial fission similarly in both cell lines, demonstrating that survivin cannot stop chemically triggered mitochondrial fragmentation. Under these conditions, survivin expression did not prevent OMM depolarisation (Figure 6C).

**Figure 6.**
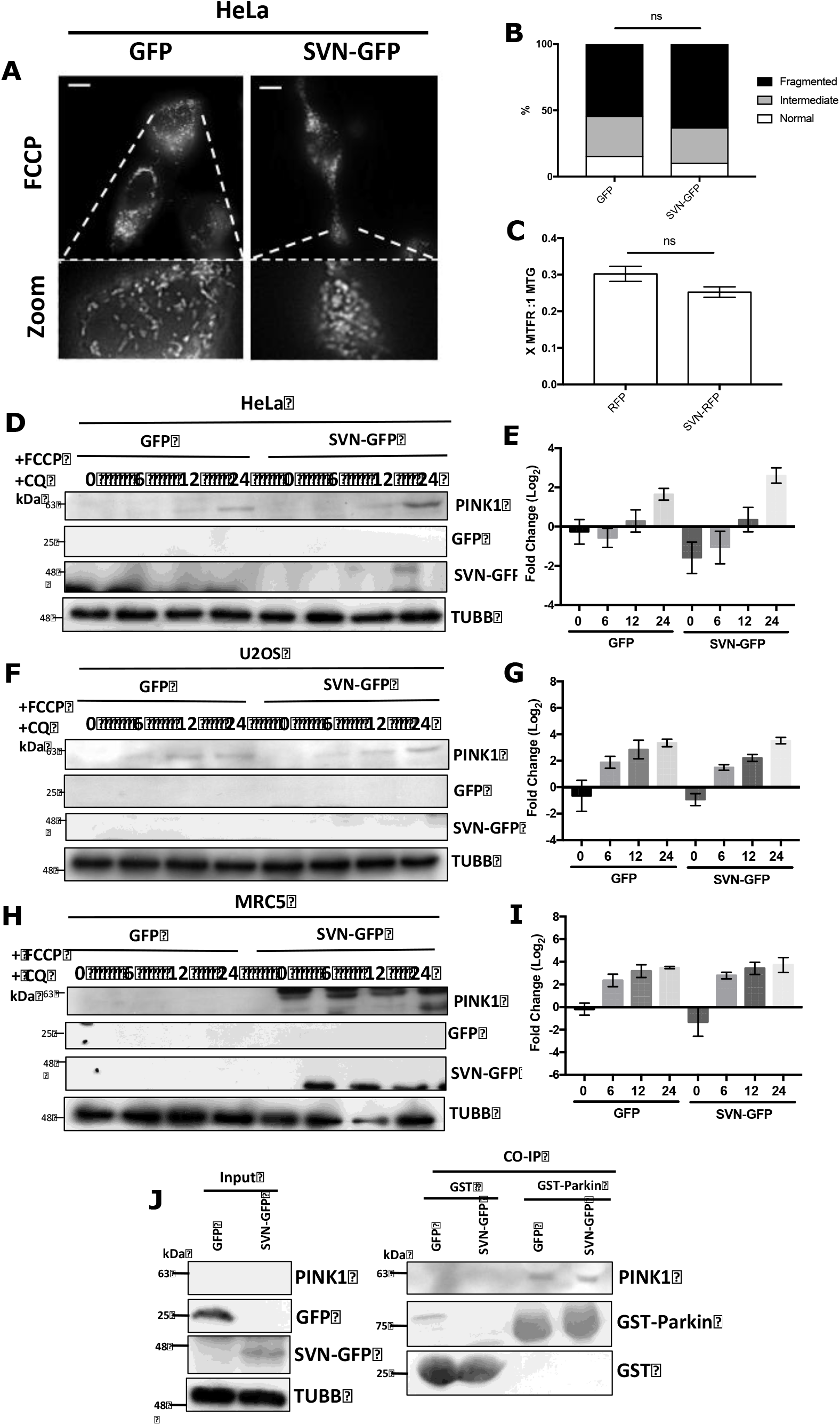
Survivin does not interfere with mitophagic stages preceding Parkin mitochondrial translocation. **(A** and **B)** HeLa cells were treated with 10 μM FCCP for 6 h, stained with 250 nM MitoTracker Red and imaged live. Mitochondrial distribution was scored as normal, fragmented, or intermediate, and no significant difference between cells expressing GFP or SVN-GFP was seen (Chi-squared test, N=3, DF=2). **(C)** HeLa cells transiently transfected with cDNA for RFP or SVN-RFP were treated as in (A) and mitochondrial membrane potential measured using MitoTracker far red, and expressed as a ratio to MTG. SVN-GFP expression did not alter MitoTracker Deep Red signal, indicating no alterations to mitochondrial membrane potential. (**D, F** and **H)** HeLa, U2OS and MRC5 cells were treated with 10 μM FCCP for 6, 12 and 24 hrs, WCEs prepared and analysed by immunoblotting. Membranes were probed for expression of the kinase PINK1, GFP to confirm cell line expression and tubulin used as a loading control N=3 (with internal triplicates). SVN-GFP expression does not alter PINK1 stabilisation post mitophagy stimulation. (**E, G** and **I)** Semi-quantification of immunoblots in **(D, F** and **H)** respectively, presented as fold change (Log2 scale) compared to the GFP control time 0. Error bars represent +/- SEM N=3 DF=32 (with internal triplicates). No statistical alterations are observed (TWO-way ANOVA). **(J)** GST-pull down assay of purified GST-Parkin and GST alone using WCE made from HeLa GFP or SVN-GFP cells. Samples were analysed by immunoblotting, and membranes probed for GST to confirm pulldown, GFP to check expression in WCE’s, and PINK1 to assess success of pulldown. Tubulin was used as a loading control. SVN-GFP does not prevent the interaction of GST-Parkin with PINK1. N=3.

Next, to determine whether survivin alters PINK1 stabilisation, HeLa, U2OS or MRC5 cells were treated with FCCP (10 μM) and Chloroquine (CQ; 100 μM) for 6, 12 or 24 h, WCE prepared and immunoblots probed for PINK1 to determine its stability post mitophagy stimulation (Figure 6D, F and H). Semi-quantification revealed no alterations to PINK1 stabilisation over the time course in either cell line (Figure 6E, G and I). Finally, to determine if survivin alters the interaction of PINK1 and its target E3-ubiquitin ligase Parkin, we performed a pulldown with recombinantly expressed GST-Parkin in the presence of a HeLa WCE expressing GFP or survivin-GFP (Figure 6J). Immunoblotting of the GST-pulldown assay shows survivin does not alter the interaction of PINK1 and GST-Parkin.

### Survivin prevents Parkin recruitment to the mitochondria

We then asked if survivin altered the recruitment of Parkin to the mitochondria post mitophagy stimulation. HeLa and MRC5 cells were transiently transfected with the mitophagy-specific E3-ligase, Parkin, N-terminally tagged with mCherry (a gift from Prof. S. Martin, Dublin), and the FCCP experiment repeated. Here a marked difference was seen: mCherry-Parkin was recruited to the mitochondria of GFP-HeLa cells after FCCP treatment, but it was retained in the cytoplasm in survivin-GFP cells (Figure 7A), and trend was confirmed by phenotype counting and quantification (Figure 7B). Conversely, the mitochondrial recruitment of mCherry-Parkin was unaffected in MRC5 cells expressing survivin-GFP (Figure 7C; images shown in Figure S5). From this we conclude that in cancer cells, after mitochondrial fragmentation and depolarisation, survivin prevents Parkin recruitment from the cytosol to the mitochondria, which blocks mitophagy.

**Figure 7.**
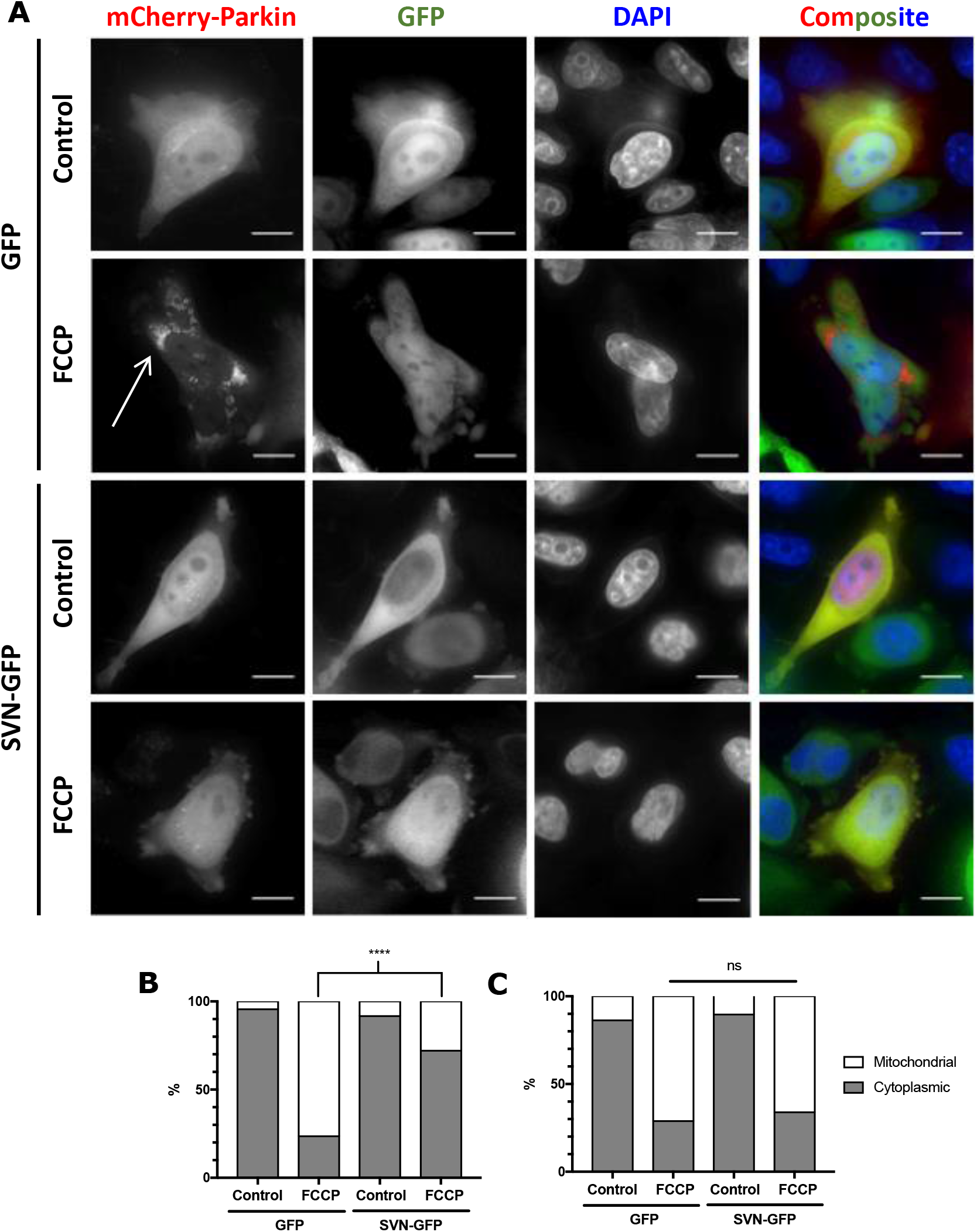
Survivin prevents mitochondrial recruitment of mCherry-Parkin. (**A**) HeLa or **(C)** MRC5 cells were treated with 10 μM FCCP post-transfection with cDNA encoding mCherry-Parkin and stained with NucBlue to visualise nuclei. Representative images, thresholded using Fiji software, scale bars 15μm. (**B** and **C**) Cells were counted for mitochondrial or cytoplasmic localisation of mCherry-Parkin, and a Chi-squared test performed to analyse differences in phenotypes. (**B)** In HeLa cells, Parkin relocates to the mitochondria in GFP cells treated with FCCP but remains cytoplasmic in SVN-GFP cells. (P-value ****<0.0001, N = 3). (**C**) No alterations were observed in mCherry-Parkin translocation in MRC5 cells (P-value 0.6748 and 0.4112, N=3).

### Survivin decreases mitochondrial co-localisation with lysosomes after mitophagy stimulation

To further confirm how survivin affects mitophagy, HeLa cells expressing RFP or SVN-RFP were treated with FCCP as previously described, and stained with LysoTracker Blue to observe autophagosomes, and MitoTracker Green to observe mitochondria (Figure 8A, S6). Co-localisation analysis was then carried out to assess the proportion of the mitochondrial network that co-localised with lysosomes. This demonstrated that post FCCP treatment, more mitochondria and lysosomes co-localised in RFP-expressing cells than in SVN-RFP cells, as shown by pixel intensity line plots (Figure 8B and C) and whole pixel co-localisation analysis of images (Figure 8D).

**Figure 8.**
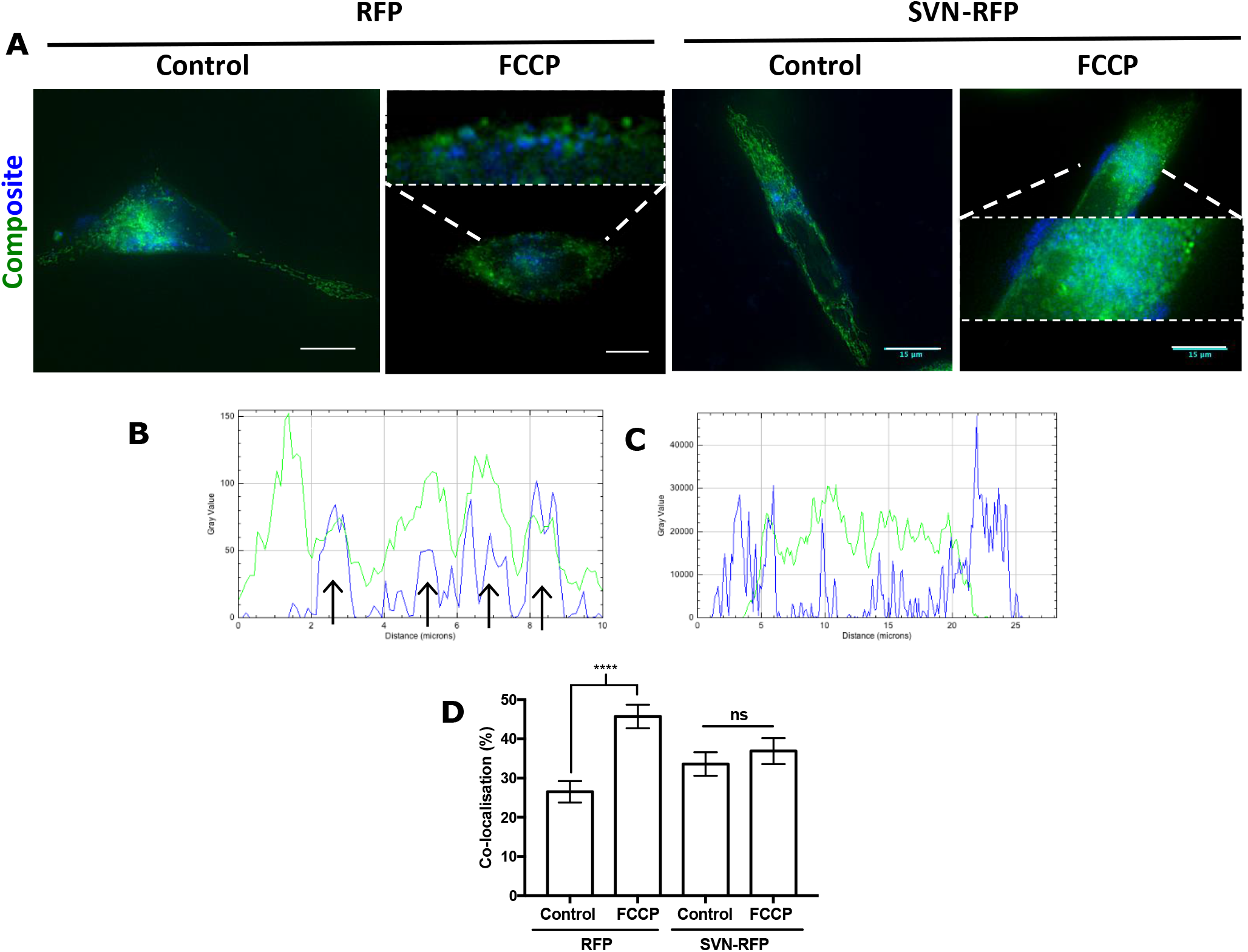
Mitochondrial co-localisation with lysosomes is reduced in survivin-RFP expressing HeLa cells. (**A**) HeLa cells were treated with 10 μM FCCP post-transfection with cDNA encoding RFP or survivin-RFP and stained with 75nM LysoTracker Blue to visualise lysosomes and 250nM MitoTracker Green for mitochondria. Representative images, thresholded using Fiji Software, scale bars 15 μm. Expanded sections to show co-localisation of mitochondria and lysosomes. (**B** and **C**) Pixel intensity profile plots over a line draw through a section of RFP (**B**) and survivin-RFP (**C**) cells treated with FCCP. RFP cells show a co-localisation of LysoTracker Blue and MitoTracker Green peaks. (**D**) Fiji co-localisation analysis of MitoTracker Green and LysoTracker Blue pixels, shown as percentage co-localisation. ONE-way ANOVA analysis shows significant increase in co-localisation after FCCP treatment of RFP cells, but no alteration to survivin-RFP cells (P-value ****<0.0001, N=3). Error bars mean +/- SEM.

### Survivin mimics the effect of Bcl-2 upon mitophagy

Finally, to determine whether of survivin collaborates with the apoptosis inhibitor Bcl-2 during mitophagy, HeLa cells expressing GFP or survivin-GFP were treated with the Bcl-2 inhibitor Navitoclax (1μM) and 10 μM FCCP to stimulate mitophagy post transfection with mCherry-Parkin. Here, mCherry-Parkin translocation to the mitochondrion was increased after FCCP treatment with Navitoclax in cells expressing survivin-GFP, but not in GFP expressing cells (Figure S7A) quantified in (Figure S7B). A UV dose response curve was simultaneously performed to prove that Navitoclax (1μM) was sufficient to inhibit Bcl-2 activity (Figure S7C). From these data we conclude the effect of survivin on mitophagy may be increased by collaboration with Bcl-2.

## Discussion

Survivin is an essential protein that is deregulated in cancer, becoming present throughout the cell cycle, rather than being confined to G2 and M-phases (Barrett, Osborne and Wheatley, 2009). In transformed cells in interphase it is predominantly cytoplasmic, shuttling between the cytoplasm and nucleus in a CRM1/exportin-dependent manner (Colnaghi *et al*., 2006; Engelsma *et al*., 2007; Stauber, Mann and Knauer, 2007). Cytoplasmic survivin inhibits apoptosis, and it has been suggested that prior residence in the mitochondria can enhance this activity (Dohi, Xia and Altieri, 2007). Consistent with previous studies (Dohi *et al*., 2004; Dohi, Xia and Altieri, 2007; Hagenbuchner *et al*., 2013, 2016; Rivadeneira *et al*., 2015), we have found that survivin only accesses the mitochondria of transformed cells. Although survivin is essential, presumably its mitochondrial residence is not, and constitutes a gain of function over its normal roles.

In this study we tested the hypothesis that survivin interferes with mitochondrial homeostasis and alters respiratory dependence in cancer cells. Mitochondria are dynamic organelles that regulate cellular metabolism and survival. The opposing pathways of mitochondrial biogenesis and autophagic degradation control their quantity and quality. Combined with fusion and fission, these mechanisms govern mitochondrial activity (Palikaras, Lionaki and Tavernarakis, 2015), and alterations to any one of these processes have been linked to ageing and disease (Redmann *et al*., 2014). In cancer cells these processes are often deregulated, consequently mitochondrial health is compromised: mtDNA harbouring mutations accumulate, respiratory efficiency declines, and ultimately cells switch from OXPHOS to glycolytic dependence (Merz and Westermann, 2009) (Sumpter *et al*., 2016). OXPHOS itself plays a major role in mtDNA damage as it produces ROS that continuously bombard the mtDNA causing lesions (Ray, Huang and Tsuji, 2012)(Sabharwal and Schumacker, 2014). Healthy cells respond to this damage by removing the affected sections of mitochondria using a selective form of autophagy called “mitophagy” (see Figure S8).

Mitophagy commences with mitochondrial fission, which produces asymmetrical daughter mitochondria, one with an increased membrane potential that can fuse with healthy mitochondria (Twig *et al*., 2008), and one with a depolarised membrane that is targeted for mitophagy (Elmore *et al*., 2001; Nicholls, 2004). Depolarisation of the OMM of defunct mitochondria stabilises the serine/threonine kinase PINK1, which phosphorylates the E3-ligase Parkin at Ser65, and activates it. Parkin then accumulates at the OMM (Vives-Bauza *et al*., 2010; Youle and Narendra, 2011) where it mediates ubiquitination of VDAC1 (Kazlauskaite *et al*., 2014). In turn VDAC-ubiquitination stimulates translocation of the autophagic adaptor protein, p62 to the mitochondria, which signals their engulfment by pre-autophagosomes via interaction with LC3 (Lee *et al*., 2010; East and Campanella, 2016).

As indicated by increased expression of VDAC, mtDNA copy number and MitoTracker Green pixel area, ectopic expression of survivin caused an increase in total mitochondrial mass. We also noted that VDAC was being affected post-translation, while FIS1 levels, which decreased when survivin levels were elevated, were affected by transcriptional repression. We have also found that survivin does not increase mitochondrial biogenesis, which combined with the aforementioned data, suggests survivin specifically increases mitochondrial mass by inhibiting mitophagy. This is further confirmed by the reduction in co-localisation of lysosomes and mitochondria post mitophagy stimulation.

In addition to the increase in mitomass, we found that cancerous cells expressing survivin had poor quality mtDNA, and that survivin suppressed OXPHOS and increased respiratory dependency on glycolysis in these cells. In general, depleting survivin had the converse effect to overexpression, and all effects were specific to cancer cells. Moreover, using a *bona fide* mitochondrial-targeting signal from cytochrome c oxidase subunit VIIIA we were able to force non-cancerous MRC5 cells to acquire the same mitochondrial characteristics as the cancerous cells tested. It is well documented that when mitophagy is inhibited, mitochondria with mtDNA lesions accumulate within the cell. This can directly impact the activity of mtDNA encoded proteins, notably members of the electron transport chain (Sumpter *et al*., 2016) which directly impacts mitochondrial metabolism. Mitophagy can be artificially blocked using general autophagy inhibitors, such as CQ, which has been shown to reduce mitochondrial metabolism in the described manner (Redmann *et al*., 2017a). Here, we saw no further reduction in the respiration of mitochondria isolated from survivin-GFP expressing cancer cells after treatment with CQ (Figure 4), therefore allowing us to conclude that survivin modifies mitochondrial metabolism specifically due to alterations to mitophagy. Survivin inhibits mitophagy, causing an accumulation of respiratory defective organelles which in turn reduces OXPHOS (Redmann *et al*., 2017b). Moreover, as OXPHOS was altered to the same degree by CQ treatment in MRC5 cells irrespective of survivin expression, we conclude that this change is due to the mitochondrial pool of survivin.

When determining where survivin acts in the mitophagic pathway, we found that neither mitochondrial fission, OMM depolarisation, PINK1 stabilisation nor its interaction with the E3 ubiquitin ligase Parkin was affected. Instead survivin prevents the recruitment of Parkin to the mitochondria. It has previously been reported that survivin inhibits the activities of PINK/Parkin, and that this response causes survivin degradation (Hagenbuchner *et al*., 2016). While our study also links survivin and Parkin, we offer a slightly alternative interpretation: rather than causing its own demise, survivin inhibits mitophagy by preventing Parkin from translocating to the mitochondria, and the resulting accumulation of mitochondria with damaged mtDNA ultimately forces the cell to switch from oxidative phosphorylation to glycolysis. As we have previously shown that survivin can inhibit autophagic flux (Humphry and Wheatley, 2018), we suggest that survivin can interfere with mitophagy both by inhibiting Parkin recruitment to the OMM, and later, preventing flux (Figure S8). Although a less efficient means of respiration, glycolysis produces less ROS, and thus, in addition to the initial survival response, this switch can provide cancer cells with a further survival advantage.

Our findings with survivin mirror these described by (Hollville *et al*., 2014a), in a series of experiments examining the role of the apoptosis inhibitor BCL-2. Accordingly, treatment with the BCL-2 inhibitor Navitoclax partially recovered the translocation of Parkin to the mitochondria, suggesting survivin acts in this process through a collaboration with BCL-2.

Finally, as none of these changes occurred in normal fibroblasts in which survivin is not mitochondrial, and in fact forcing survivin into the mitochondria of these cells decreases metabolism, we conclude that the effects are exclusive to cancer cells and can be attributed solely to the mitochondrial pool of survivin. The targeting of mitochondrial survivin and the metabolic alterations it provides cancerous cells, could therefore offer a distinct opportunity to develop novel therapeutic treatments with reduced off target effects in non-cancerous cell lines.

## Materials and Methods

All reagents were obtained from Sigma unless specified.

### Human Cell Culture

Human epithelial carcinoma cells (HeLa, ATCC), Osteosarcoma (U2OS), and normal lung fibroblasts (MRC5:Medical Research Council Strain 5, Genome Stability Centre, Sussex), were cultured in 5% CO_2_ at 37°C with humidity in Dulbecco’s Modified Eagle’s Medium (DMEM, Gibco, Invitrogen) supplemented with 4 mM L-glutamine, 10% Fetal Calf Serum (FCS, Thermo Scientific), 244 μM penicillin and 172 μM streptomycin. Derivative lines expressing GFP or survivin-GFP were maintained under the selection pressure of 1 mM G418 (Fisher). Experiments were carried out on cells within 30 passages.

### DNA and siRNA Transfections

Cells were seeded into a relevant dish or imaging chamber in antibiotic free media, incubated for 12 h before transfection and left for approximately 48 h before use. DNA transfections were performed using Torpedo Transfection reagent (Ibidi) and 0.3 μg of relevant DNA construct, as per the manufacturer’s guidelines. siRNA transfections were performed using HiPerfect transfection reagent (Qiagen) and 75 ng of relevant siRNA, as per the manufacturer’s instruction.

### DNA constructs

| Construct | Vector | Reference |
| --- | --- | --- |
| GFP | pcDNA 3.1+ | (Carvalho *et al.*, 2003) |
| Survivin-GFP | pcDNA3.1+ | (Carvalho *et al.*, 2003) |
| MTS-survivin-GFP | pcDNA3.1+ | MTS of cytochrome c oxidase was amplified by PCR from U2OS cDNA with EcoR1 cut sites and inserted into SVN-GFP pcDNA3.1+ |
| RFP | pRFP1-N1 | Clontech |
| Survivin-RFP | pRFP1-N1 | Survivin was cut from survivin-GFP pcDNA3.1+ using HindIII and inserted into pRFP-N1. |
| mCherry-Parkin | mCherry-C1 | mCherry-Parkin was a gift from Seamus J. Martin (Hollville *et al.*, 2014b) |
| GST | pGEX4T1 | Amersham. |
| GST-Parkin-WT | pGEX4T1 | pGEX-parkin WT was a gift from Kalle Gehring (Addgene) |
|  |  | plasmid #45969;<br>http://n2t.net/addgene:45969;<br>RRID:Addgene_45969)<br>(Trempe *et al.*, 2013) |

### Cell counting assay

Cells were seeded at a density of 200 cells per 10 cm petri dish, and the number of cells in individual colonies after 8, 24, 48 and 72 h.

### Immunoblotting

Protein samples were separated according to standard procedures using 12% acrylamide SDS-PAGE gels, in running buffer (25 mM Tris, 192 mM glycine, 0.1% (w/v) SDS) and transfer onto a 0.22 μM nitrocellulose membrane (BIOTRACE, PALL life sciences) using transfer buffer (24 mM Tris, 195 mM glycine, 0.1% SDS, 10% methanol). Post-transfer membranes were blocked with 5% non-fat milk (Marvel, in PBS +0.1% Tween 20 (PBST)) then incubated with appropriate primary antibody overnight at 4°C, washed three times with PBST then incubated in the appropriate horseradish peroxidase conjugated secondary antibody (see 3.5) in 5% non-fat milk. EZ-ECL chemiluminescence detection reagent was then added (Geneflow) and membrane exposed to detection film (Roche).

Immunoblots were quantified using Fiji software. Band intensity peaks were measured and combined into sample groups for each condition. Within each pool, intensity values for each protein were expressed as a percentage of the loading control average and then as a percentage of the control protein average. The final expression value was presented as a decimal and transformed as a function of a base 2 logarithm (log2).

### Mitochondrial DNA lesion assay

2 x 10^6^ cells were washed with PBS and harvested by scraping. Genomic DNA was extracted using the GeneJET genomic DNA purification Kit (Thermo Scientific #K0721) according to the manufacturer’s instructions.

To determine mitochondrial DNA integrity, a PCR reaction was performed on gDNA samples using 6 ng of template gDNA, 250 μM dNTPs, 500 nM of either a short read (tRNA(LEU)) or a long read (LR-mtDNA) primer mix, 0.02U/μl Q5 DNA polymerase and 1X Q5 reaction buffer. Long read PCR was aided by the addition of 10 ng/μl BSA. PCR products were then run on a 0.8% agarose gel and quantified using Fiji software as described for immunoblots.

### RNA extraction

7 x10^6^ cells were harvested by scraping into 0.2 ml of media, 1 ml TRI-reagent was added, and the samples incubated (5 minutes at RT). 200 μl 1-Bromo-3-chloropropane (BCP; 11.76% (v/v) in TRI reagent and residual DMEM) was then added and the sample incubated for a further 3 minutes at RT before centrifugation (10,000 x g, Labnet, Prism R). The upper (colourless) layer was removed and incubated overnight at −20°C in acidified isopropanol solution (256 mM sodium acetate pH 4 and 36% isopropanol (v/v)). Samples were centrifuged at 17,000 x g for 15 minutes at 4°C and pellets washed in 70% ethanol. Samples were DNase treated with RNAse free DNase-I (Qiagen) according to the manufacturer’s instructions. RNA was isolated by phenol-chloroform-isoamyl-alchohol (50% phenol-chloroform-isoamy alchohol (v/v) 125:24:1 in dH_2_O. Samples were vigorously shaken and incubated for 3 minutes at RT before centrifugation. The upper aqueous phase was transferred to a fresh tube and incubated overnight at −20 °C in acidified ethanol solution (323mM sodium acetate pH 5.2 and 65% ethanol (v/v)). Precipitated samples were washed in 70% ethanol, pellets dried, dissolved in dH_2_O and RNA concentration determined by spectrophotometry (Nanodrop 2000, Thermo Scientific).

### Comparative qPCR

qPCR was performed using iTaq^™^ Universal SYBR^®^ Green Supermix (BIORAD) as per the manufacturer’s instructions on a qPCR thermocycler (7500 fast real-time qPCR, Applied Biosciences). gDNA samples were analysed at a final concentration of 200 pg/μl with 500 nM primers. Data sets were analysed, and reference genes verified using the Pfaffl method and REST software (QIAGEN).

### Subcellular fractionation

Cells grown to 80-90% confluence in 15 cm^2^ petri dishes were washed and scraped into PBS and pelleted at 300 g for 3 minutes, before re-suspension in 2ml homogenisation buffer (200 mM Mannitol, 70 mM Sucrose, 1 mM EGTA, 10 mM HEPES, pH 7.5) supplemented with protease and kinase inhibitors. Lysates were prepared in a glass homogeniser (Teflon), a sample taken as whole cell extract, before spinning at 1000 g, 4°C for 5 minutes. Supernatants were transferred to a fresh tube (mitochondrial/cytoplasmic fraction) and centrifuged at 10,000 g, 4°C for 15 minutes to pellet mitochondria before re-suspension in homogenisation buffer. Protein concentration was then measured by Bradford assay and samples were boiled in SDS sample buffer, and 20 μg protein loaded onto an SDS-page gel for analysis.

### Resazurin assays

Mitochondrial metabolism assays were performed using 20 μg of isolated mitochondria were re-suspended in Locke’s buffer (154 mM NaCl, 5.6 mM KCl (BDH), 2.3 mM CaCl_2_ (Fischer), 1 mM MgCl_2_, 3.6 mM NaHCO_3_, 5 mM Glucose and 5 mM HEPES pH 7.5 (BDH) and plated into 1 well of a black 96 well plate (CLS3904). 40 μM resazurin was then added to each sample to make a final concentration of 20 μM and plates were then incubated at 37°C with 5% CO_2_. Absorbance was then read at 595nm every 30 minutes for 4 h and compared to that of 20 μM resazurin blank controls (FluoStar Galaxy).

Cell death curves were performed on 50,000 HeLa cells expressing GFP, pre-treated with and without 1 μM of the BCL-2 inhibitor Navitoclax for 16h. Cells were treated with 0, 0.02, 0.08, 0.32, 1.28 or 2.56 J of UV, left for 24 h, 20 μM of resazurin added and plates incubated at 37°C with 5% CO_2_ for 1h. Absorbance was read at 595nm and compared to 20 μM resazurin control.

### Glucose-Glo and Lactate-Glo assay

10,000 HeLa cells were plated per well of a 96 well plate in 100 μl DMEM (Gibco 11054001) containing 5.6 mM glucose, 2 mM glutamine and supplemented with 10% FCS. Media only wells acted as controls. At 8, 24, 48 and 72 h post plating, 2.5 μl of media was removed from each sample, diluted in 97.5 μl of PBS and frozen until needed. On day of assay, samples thawed and were diluted a further 2.5 x, and 50 μl of diluted media added to a white 96 well plate (CLS3610) before the addition of 50 μl of Lactate/Glucose detection reagent (Promega). Plates were incubated for 1 h at RT and luminescence recorded (Glowmax Luminometer, Promega) and then compared to Glucose/Lactate standards to determine relevant concentrations.

### Live Cell Fluorescence imaging

To visualise active mitochondria, cells were grown overnight in 8-chambered micro-slides (Ibidi). Cells were stained to visualise mitochondria using either 500 nM MitoTracker Red CMXRos, MitoTracker Deep Red FM or 200nM MitoTracker Green FM (Invitrogen), and Nucblue to visualise DNA (Thermo Fischer Scientific) in phenol-red free CO_2_ independent media (DMEM, Invitrogen) supplemented with 2 mM L-glutamine and 10% Fetal Calf Serum, for 15 minutes at 37°C.

### Mitochondrial membrane potential assay

HeLa or MRC5 cells were seeded into Ibidi 8-chambered chambers approximately 24 h before imaging and once adherent, were incubated in DMEM without phenol red (D1145) supplemented with 4 mM L-glutamine and 25 mM HEPES. 15 minutes before imaging, cells were stained with 1 drop of Nucblue (Life Technologies), 100 nM MitoTracker Green FM and 100 nM MitoTracker Deep Red FM or 200 μM tetramethylrhodamine ethyl ester perchlorate (TMRE, AAT Bioquest). Immediately before imaging, stains were washed off by replacing media with phenol red free CO_2_–independent complete DMEM.

### Image acquisition and processing

Imaging was performed using an inverted (DMRIB Olympus, Delta Vision Elite) microscope with a 60x (NA1.4, oil) objective. Single plane images were acquired, de-convolved using inbuilt software on the Delta-vision and saved as TIFF files. Image pixel intensity was quantified in Fiji using a fully automated macro, which was programmed first to threshold each channel to a set scale defined by the user, and then measure the average signal intensity of each channel. Co-localisation analysis was performed using a similar macro that after thresholding images, was then programmed to analyse the percentages of pixels that spatially co-localised with similar intensity between the two channels. Datasets were then analysed in the GraphPad Prism software.

## Supplementary Data

**Table 1:**
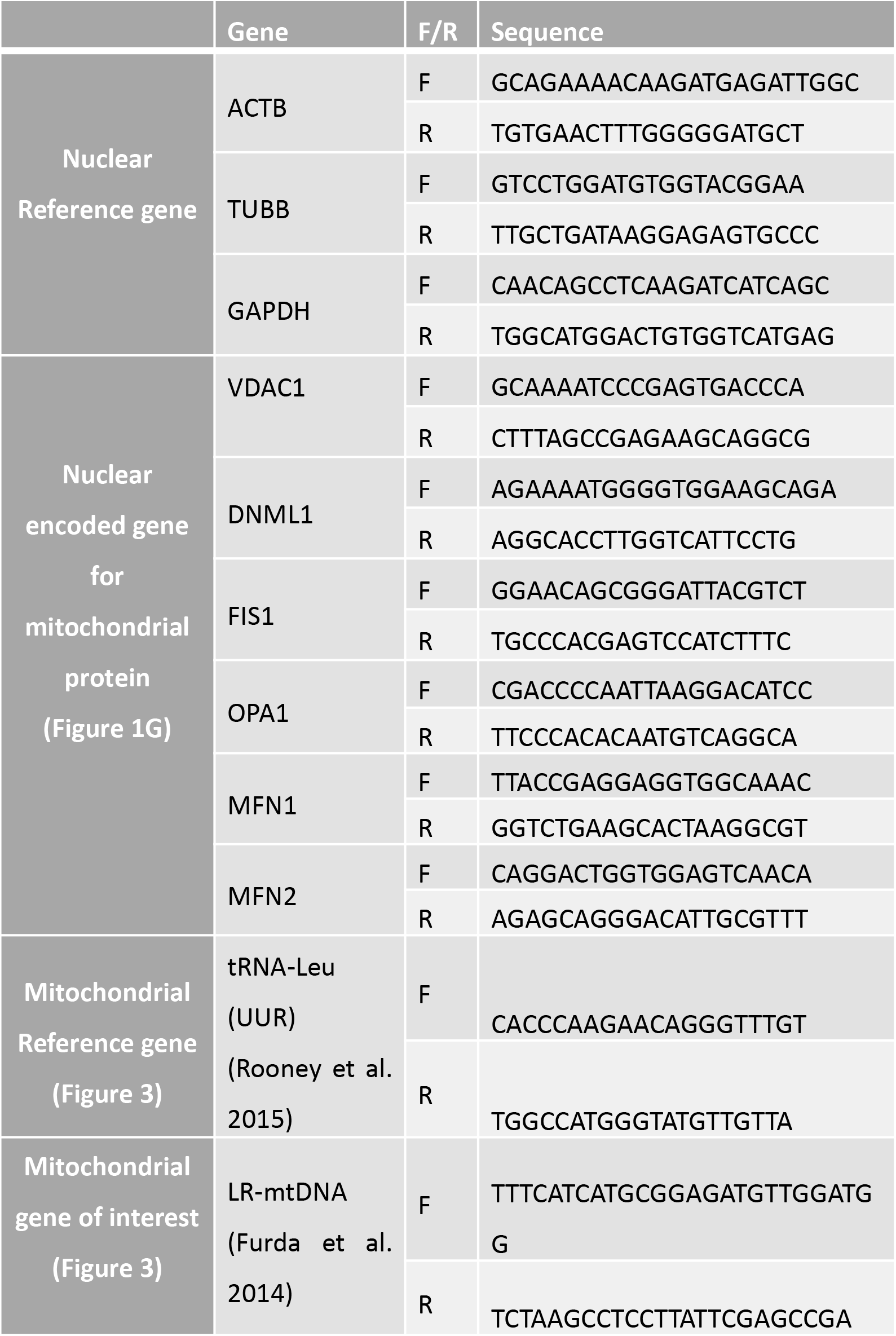
**qPCR primers,** primer sequences for nuclear reference genes and mitochondrial genes of interest. F: forward; R: reverse.

***Figure S1. Survivin up-regulation increases mitomass but does not by altered mitochondrial biogenesis*. (A)** Hela, or **(B)** MRC5 cells were treated with mock siRNA or survivin siRNA, WCEs prepared and analysed by immunoblotting. Membranes were probed for expression of survivin, the mitochondrial proteins, VDAC, MFN1, FIS1, OPA1, and DRP1, and tubulin used as a loading control N=2 (with internal triplicates). **(C)** and **(D)** Semi-quantification of immunoblots in **(A)** and **(B)** respectively, presented as fold change (Log2 scale) compared to the GFP control. (TWO-way ANOVA, * p<0.05, ** p<0.01, **** p<0.0001). Error bars represent +/- SEM N=2 (with internal triplicates). Survivin depletion reduces the expression of VDAC and MFN1.

***Figure S2*.** Each MRC5 data set shown in Figure 3E presented individually for clarity.

***Figure S3. Mitochondrial fragmentation is unaffected by survivin-GFP expression***. Full figure to accompany Figure 5A. Scale bars 10 μm.

***Figure S4. Mitochondrial polarisation (TMRE) and distribution (MitoTracker green) after survivin siRNA*. (A)** HeLa and **(B)** MRC5 cells. Full figure accompanying Figure 5E-I. Scale bars 10 μm.

***Figure S5. (A) mCherry-Parkin mitochondrial recruitment in MRC5 cells*.** Full figure accompanying Figure 7C. Scale bars 10 and 15 μm respectively. ***(B) Inhibition of Bcl-2 increases mitochondrial translocation of Parkin in survivin-GFP expressing HeLa cells*.** HeLa cells expressing GFP or survivin-GFP were treated with 10 μM FCCP and 1 μM Navitoclax post-transfection with cDNA encoding mCherry-Parkin. Cells were counted for mitochondrial or cytoplasmic localisation of mCherry-Parkin, and a Chi-squared test performed to analyse differences in phenotypes. Images accompany Figure 10E.

***Figure S6*.** Full figure accompanying Figure 8A. Scale bars 15 μm.

***Figure S7. Inhibition of BCL-2 increases mitochondrial translocation of Parkin in survivin-GFP expressing HeLa cells*.** (**A**) HeLa cells expressing GFP or SVN-GFP were treated with 10 μM FCCP and 1 μM Navitoclax post-transfection with cDNA encoding mCherry-Parkin. (**B**) Cells were counted for mitochondrial or cytoplasmic localisation of mCherry-Parkin, and a binomial test performed to test for differences to expected verus observed phenotypes. BCL-2 inhibition in survivin up-regulated HeLa cells caused increased mCherry-Parkin mitochondrial recruitment post FCCP treatment (****p=<0.0001). N=3, error bars represent mean +/- SEM. (**C**) Resazurin assay performed on 50,000 HeLa cells expressing GFP treated +/- 1 μM Navitoclax and increasing concentrations of UVC. Cells were left for 24 h post UV treatment, 20 μM resazurin added and absorbance read to determine cell survival. N=1 (with internal triplicates), error bars mean +/- SEM.

***Figure S8. Working model: Survivin inhibits Parkin-mediated mitophagy*.** Healthy mitochondria have polarized membranes and respire by OXPHOS. On encountering stress their mtDNA is damaged and their health compromised. Cells deal with this damage by fragmenting their mitochondria asymmetrically to generate two derivatives, one healthy mitochondrion and one that contains the defective components, which is then targeted for removal from the cell by a specialized form of autophagy, called mitophagy. Fragmentation, also called “fission” involves two proteins, FIS1 and DRP1, and occurs simultaneously with the depolarization of the OMM of the defective mitochondrion. Subsequently, the mitophagy specific E3-ubiquitin ligase, Parkin, phosphorylated by PINK, is recruited to the OMM. In the absence of survivin the OMM proteins, VDAC and MFN1 get polyubiquitinated, which signals to adaptor proteins, such as p62, to build a pre-autophagosome around the defective organelle. Extension of the pre-autophagosomal membrane, which is facilitated by LC3II, then enables the total engulfment of the defective mitochondria. Finally, the autophagosome fuses with a lysosome thus completing the clearance of defunct mitochondria from the cell. The data herein presented, suggest that survivin blocks mitophagy by preventing recruitment of Parkin to the OMM. Thus, we propose that by inhibiting mitophagy, survivin causes an accumulation of defunct mitochondria, which causes cancer cells to switch their respiratory dependency from OXPHOS to glycolysis, aka the “Warburg Effect”.

## Acknowledgements

We thank Dr. Sophie Rochette for technical support, Alex Fezovich for assistance with FiJi and Prof. Seamus Martin for mcherry-PARKIN. A.Townley is a BBSRC-DTP funded student, we thank the BBSRC for her support.

## References

Ambrosini, G., Adida, C. and Altieri, D. C. (1997) ‘A novel anti-apoptosis gene, survivin, expressed in cancer and lymphoma’, Nature Medicine. 3(8), pp. 917–921. doi: 10.1038/nm0897-917.

Balaban, R. S., Nemoto, S. and Finkel, T. (2005) ‘Mitochondria, Oxidants, and Aging’, Cell, 120(4), pp. 483–495. doi: 10.1016/j.cell.2005.02.001.

Barrett, R. M. A., Osborne, T. P. and Wheatley, S. P. (2009) ‘Phosphorylation of survivin at threonine 34 inhibits its mitotic function and enhances its cytoprotective activity’, Cell Cycle, 8(2), pp. 278–283. doi: 10.4161/cc.8.2.7587.

Carvalho, A. et al. (2003) ‘Survivin is required for stable checkpoint activation in taxol-treated HeLa cells.’, Journal of Cell Science. 116(Pt 14), pp. 2987–98. doi: 10.1242/jcs.00612.

Chatterjee, A., Dasgupta, S. and Sidransky, D. (2011) ‘Mitochondrial subversion in cancer.’ 4(5), pp. 638–54. doi: 10.1158/1940-6207.CAPR-10-0326.

Colnaghi, R. et al. (2006) ‘Separating the Anti-apoptotic and Mitotic Roles of Survivin’, Journal of Biological Chemistry. 281(44), pp. 33450–33456. doi: 10.1074/jbc.C600164200.

Dohi, T. et al. (2004) ‘Mitochondrial survivin inhibits apoptosis and promotes tumorigenesis’, Journal of Clinical Investigation, 114(8), pp. 1117–1127. doi: 10.1172/JCI200422222.

Dohi, T., Xia, F. and Altieri, D. C. (2007) ‘Compartmentalized phosphorylation of IAP by protein kinase A regulates cytoprotection.’, Molecular cell. 27(1), pp. 17–28. doi: 10.1016/j.molcel.2007.06.004.

East, D. A. and Campanella, M. (2016) ‘Mitophagy and the therapeutic clearance of damaged mitochondria for neuroprotection’, The International Journal of Biochemistry & Cell Biology, 79, pp. 382–387. doi: 10.1016/j.biocel.2016.08.019.

Elmore, S. P. et al. (2001) ‘The mitochondrial permeability transition initiates autophagy in rat hepatocytes.’, FASEB journal, 15(12), pp. 2286–7. doi: 10.1096/fj.01-0206fje.

Engelsma, D. et al. (2007) ‘Homodimerization Antagonizes Nuclear Export of Survivin’, Traffic. 8(11), pp. 1495–1502. doi: 10.1111/j.1600-0854.2007.00629.x.

Escuín, D. and Rosell, R. (1999) ‘The Anti-Apoptosis Survivin Gene and its Role in Human Cancer: An Overview’, Clinical Lung Cancer. 1(2), pp. 138–143. doi: 10.3816/CLC.1999.n.011.

Fortugno, P. et al. (2002) ‘Survivin exists in immunochemically distinct subcellular pools and is involved in spindle microtubule function.’, Journal of cell science, 115(Pt 3), pp. 575–85.

van Gisbergen, M. W. et al. (2015) ‘How do changes in the mtDNA and mitochondrial dysfunction influence cancer and cancer therapy? Challenges, opportunities and models’, Mutation Research/Reviews in Mutation Research. 764, pp. 16–30. doi: 10.1016/J.MRREV.2015.01.001.

Hagenbuchner, J. et al. (2013) ‘BIRC5/Survivin enhances aerobic glycolysis and drug resistance by altered regulation of the mitochondrial fusion/fission machinery.’, Oncogene. 32(40), pp. 4748–57. doi: 10.1038/onc.2012.500.

Hagenbuchner, J. et al. (2016) ‘BIRC5/Survivin as a target for glycolysis inhibition in high-stage neuroblastoma’, Oncogene, 35, pp. 2052–2061. doi: 10.1038/onc.2015.264.

Hollville, E. et al. (2014) ‘Bcl-2 Family Proteins Participate in Mitochondrial Quality Control by Regulating Parkin/PINK1-Dependent Mitophagy’, Molecular Cell, 55(3), pp. 451–466. doi: 10.1016/j.molcel.2014.06.001.

Humphry, N. J. and Wheatley, S. P. (2018) ‘Survivin inhibits excessive autophagy in cancer cells but does so independently of its interaction with LC3.’, Biology Open. 7(10), p. bio037374. doi: 10.1242/bio.037374.

Jaiswal, P. K., Goel, A. and Mittal, R. D. (2015) ‘Survivin: A molecular biomarker in cancer.’, The Indian journal of medical research. 141(4), pp. 389–97. doi: 10.4103/0971-5916.159250.

Jornayvaz, F. R. and Shulman, G. I. (2010) ‘Regulation of mitochondrial biogenesis.’, Essays in biochemistry. NIH Public Access, 47, pp. 69–84. doi: 10.1042/bse0470069.

Kazlauskaite, A. et al. (2014) ‘Parkin is activated by PINK1-dependent phosphorylation of ubiquitin at Ser 65’, Biochemical Journal, 460(1), pp. 127–141. doi: 10.1042/BJ20140334.

Lee, J.-Y. et al. (2010) ‘Disease-causing mutations in parkin impair mitochondrial ubiquitination, aggregation, and HDAC6-dependent mitophagy.’ The Journal of Cell Biology. 189(4), pp. 671–9. doi: 10.1083/jcb.201001039.

Merz, S. and Westermann, B. (2009) ‘Genome-wide deletion mutant analysis reveals genes required for respiratory growth, mitochondrial genome maintenance and mitochondrial protein synthesis in Saccharomyces cerevisiae.’, Genome biology. 10(9), p. R95. doi: 10.1186/gb-2009-10-9-r95.

Morrison, D. J. et al. (2012) ‘Endogenous knockdown of survivin improves chemotherapeutic response in ALL models.’, Leukemia. 26(2), pp. 271–9. doi: 10.1038/leu.2011.199.

Nicholls, D. G. (2004) ‘Mitochondrial membrane potential and aging.’ Aging cell, 3(1), pp. 35–40..

Nunnari, J. et al. (1997) ‘Mitochondrial transmission during mating in Saccharomyces cerevisiae is determined by mitochondrial fusion and fission and the intramitochondrial segregation of mitochondrial DNA.’, Molecular biology of the cell. American Society for Cell Biology, 8(7), pp. 1233–42.

Ott, M. et al. (2007) ‘Mitochondria, oxidative stress and cell death’, Apoptosis, 12, pp. 913–922. doi: 10.1007/s10495-007-0756-2.

Palikaras, K., Lionaki, E. and Tavernarakis, N. (2015) ‘Balancing mitochondrial biogenesis and mitophagy to maintain energy metabolism homeostasis’, Cell Death & Differentiation, 22(9), pp. 1399–1401. doi: 10.1038/cdd.2015.86.

Pfaffl, M. W. (no date) Relative quantification. Available at: https://gene-quantification.de/pfaffl-rel-quan-book-ch3.pdf

Porporato, P. E. et al. (2017) ‘Mitochondrial metabolism and cancer’, Cell Research. doi: 10.1038/cr.2017.155.

Ray, P. D., Huang, B.-W. and Tsuji, Y. (2012) ‘Reactive oxygen species (ROS) homeostasis and redox regulation in cellular signaling.’, Cellular signalling. 24(5), pp. 981–90. doi: 10.1016/j.cellsig.2012.01.008.

Redmann, M. et al. (2014) ‘Mitophagy mechanisms and role in human diseases.’, The international journal of biochemistry & cell biology. 53, pp. 127–33. doi: 10.1016/j.biocel.2014.05.010.

Redmann, M. et al. (2017) ‘Inhibition of autophagy with bafilomycin and chloroquine decreases mitochondrial quality and bioenergetic function in primary neurons.’ Redox biology. 11, pp. 73–81. doi: 10.1016/j.redox.2016.11.004.

Rivadeneira, D. B. et al. (2015) ‘Survivin promotes oxidative phosphorylation, subcellular mitochondrial repositioning, and tumor cell invasion.’, Science signaling. 8(389), p. ra80. doi: 10.1126/scisignal.aab1624.

Sabharwal, S. S. and Schumacker, P. T. (2014) ‘Mitochondrial ROS in cancer: initiators, amplifiers or an Achilles’ heel?’, Nature Reviews Cancer, 14(11), pp. 709–721. doi: 10.1038/nrc3803.

Shaid, S. et al. (2013) ‘Ubiquitination and selective autophagy.’, Cell death and differentiation. 20(1), pp. 21–30. doi: 10.1038/cdd.2012.72.

Stauber, R. H., Mann, W. and Knauer, S. K. (2007) ‘Nuclear and cytoplasmic survivin: molecular mechanism, prognostic, and therapeutic potential.’, Cancer Research. 67(13), pp. 5999–6002. doi: 10.1158/0008-5472.CAN-07-0494.

Sumpter, R. et al. (2016) ‘Fanconi Anemia Proteins Function in Mitophagy and Immunity.’, Cell. 165(4), pp. 867–81. doi: 10.1016/j.cell.2016.04.006.

Trempe, J.-F. et al. (2013) ‘Structure of parkin reveals mechanisms for ubiquitin ligase activation.’, Science. 340(6139), pp. 1451–5. doi: 10.1126/science.1237908.

Twig, G. et al. (2008) ‘Fission and selective fusion govern mitochondrial segregation and elimination by autophagy’, The EMBO Journal, 27(2), pp. 433–446. doi: 10.1038/sj.emboj.7601963.

Twig, G., Hyde, B. and Shirihai, O. S. (2008) ‘Mitochondrial fusion, fission and autophagy as a quality control axis: the bioenergetic view.’, Biochimica et biophysica acta, 1777(9), pp. 1092–7. doi: 10.1016/j.bbabio.2008.05.001.

Twig, G. and Shirihai, O. S. (2011) ‘The interplay between mitochondrial dynamics and mitophagy.’, Antioxidants & redox signaling. 14(10), pp. 1939–51. doi: 10.1089/ars.2010.3779.

Vives-Bauza, C. et al. (2010) ‘PINK1-dependent recruitment of Parkin to mitochondria in mitophagy.’, Proceedings of the National Academy of Sciences of the United States of America. National Academy of Sciences, 107(1), pp. 378–83. doi: 10.1073/pnas.0911187107.

Warburg, O. (1956) ‘On the Origin of Cancer Cells’, Science. American Association for the Advancement of Science, 123(3191), pp. 235–314. doi: 10.1126/science.123.3191.309.

Westermann, B. (2010) ‘Mitochondrial fusion and fission in cell life and death’, Nature Reviews Molecular Cell Biology. 11(12), pp. 872–884. doi: 10.1038/nrm3013.

Westermann, B. (2010b) ‘Mitochondrial fusion and fission in cell life and death’, Nature Reviews Molecular Cell Biology. Nature Publishing Group, 11(12), pp. 872–884. doi: 10.1038/nrm3013.

Wheatley, S. P. and Altieri, D. C. (2019) ‘Survivin at a glance’, Journal of Cell Science, 132(7), p. jcs223826. doi: 10.1242/jcs.223826.

Yadav, N. and Chandra, D. (2013) ‘Mitochondrial DNA mutations and breast tumorigenesis.’, Biochimica et biophysica acta. 1836(2), pp. 336–44. doi: 10.1016/j.bbcan.2013.10.002.

Youle, R. J. and Narendra, D. P. (2011) ‘Mechanisms of mitophagy’, Nature Reviews Molecular Cell Biology. 12(1), pp. 9–14. doi: 10.1038/nrm3028.

Zaffagnini, G. and Martens, S. (2016) ‘Mechanisms of Selective Autophagy.’, Journal of molecular biology. 428(9 Pt A), pp. 1714–24. doi: 10.1016/j.jmb.2016.02.004.

